# Contamination detection and microbiome exploration with GRIMER

**DOI:** 10.1101/2021.06.22.449360

**Authors:** Vitor C. Piro, Bernhard Y. Renard

**Affiliations:** Data Analytics and Computational Statistics, Hasso Plattner Insititute, Digital Engineering Faculty, University of Potsdam, 14482 Potsdam, Germany; Department of Mathematics and Computer Science, Freie Universität Berlin, Takustr. 9, 14195 Berlin, Germany

## Abstract

**Background:** Contamination detection is a important step that should be carefully considered in early stages when designing and performing microbiome studies to avoid biased outcomes. Detecting and removing true contaminants is challenging, especially in low-biomass samples or in studies lacking proper controls. Interactive visualizations and analysis platforms are crucial to better guide this step, to help to identify and detect noisy patterns that could potentially be contamination. Additionally, external evidence, like aggregation of several contamination detection methods and the use of common contaminants reported in the literature could help to discover and mitigate contamination.

**Results:** We propose GRIMER, a tool that performs automated analyses and generates a portable and interactive dashboard integrating annotation, taxonomy and metadata. It unifies several sources of evidence to help detect contamination. GRIMER is independent of quantification methods and directly analyses contingency tables to create an interactive and offline report. Reports can be created in seconds and are accessible for non-specialists, providing an intuitive set of charts to explore data distribution among observations and samples and its connections with external sources. Further, we compiled and used an extensive list of possible external contaminant taxa and common contaminants with 210 genera and 627 species reported in 22 published articles.

**Conclusion:** GRIMER enables visual data exploration and analysis, supporting contamination detection in microbiome studies. The tool and data presented are open-source and available at: https://gitlab.com/dacs-hpi/grimer.

## Introduction

Microbiome studies enable, via high-throughput sequencing, the investigation of the composition of complex microbial communities from diverse environments. Microbiome studies usually yield large amounts of raw sequences for several samples that can be analyzed with an increasing number of computational methods and databases. Standards, protocols, and best practices for designing and performing a microbiome study have been improving and changing over the years [1, 2] and the field is in constant evolution due to higher availability and reduced costs of sequencing runs as well as with the increase in number of publicly available reference sequences and computational methods.

In early stages of a standard *in silico* microbiome analysis, raw or quality-filtered sequences are classified or clustered into specific groups and quantified to generate a profile for a given environmental sample. Marker gene, whole metagenome, or metatranscriptome analysis have their own set of tools and standards which should be carefully chosen to generate reliable measurements for each sample in the study [3]. This step can be computationally intensive but reduces the large amount of data into a concise table of measurements. Alternatively, genome assembly can be performed for metagenomics samples, allowing genome-resolved analysis. Although still a complex task, gene prediction, taxonomic and functional analysis are improved with metagenome-assembled genomes, resulting in overall better measurements [4].

After measurements are obtained, hypotheses are validated through data mining and statistical analysis. This step is mostly exploratory and specific to the hypotheses and research questions pursued and the required analyses are difficult to be fully automatized. It is also very important to take in consideration the compositionality of data at this stage when working with the microbiome [5]. Several comprehensive and generalized analytical packages [6, 7, 8] and web platforms (Table 1) are available to perform a large number of microbiome analysis: basic data summaries, diversity and functional analysis, microbial interactions, differential abundance among others. Additionally, interactive tools for analytical and visual exploration are extremely helpful in this stage to better understand the data distribution and its properties and to guide further investigations to follow. In the last decade, several applications were developed with focus on visualization of microbiome data (Table 2). A comparison among many of those methods and their functionalities can be found in a recent review [9].

**Table 1:** Web resources to process, analyze and visualize microbiome data

| Name | Website | Reference |
| --- | --- | --- |
| MG-RAST | https://www.mg-rast.org/ | [10] |
| MGnify | https://www.ebi.ac.uk/metagenomics/ | [11] |
| MicrobiomeDB | https://microbiomedb.org/ | [12] |
| Nephele | https://nephele.niaid.nih.gov/ | [13] |
| Qiita | https://qiita.ucsd.edu/ | [14] |

**Table 2:**
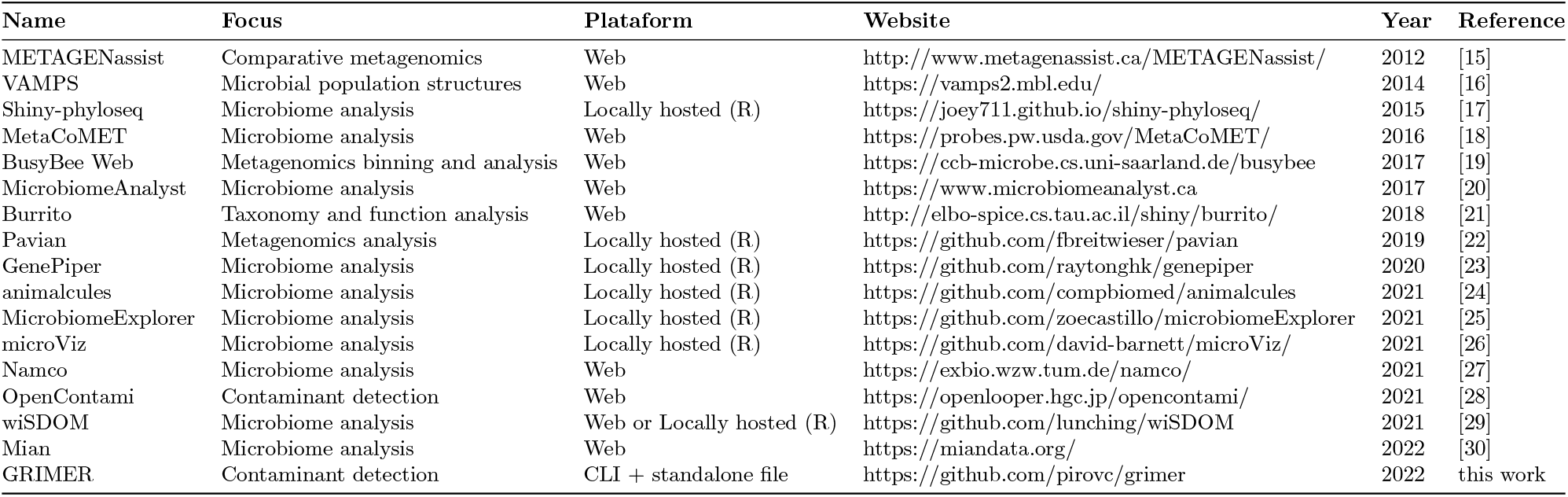
Interactive analysis and visualization tools for microbiome data published in the last 10 years

At this stage of a study, contamination detection should be considered. Contamination side-effects have gained attention in recent years due to the controversial detection of a placental microbiome [31, 32, 33]. However, the issue is not new and contamination has been known and reported for decades in the literature [34]. Contamination is characterized by exogenous DNA in a given sample introduced externally or internally. External contamination can come from diverse sources: DNA extraction kits, laboratory reagents, surfaces and equipment, ultra-pure water, residuals from previous sequencing runs as well as microbes from laboratory technicians [2, 35, 36]. Internal contamination can be defined as a undesired exchange of genetic material between samples and it is usually referred as well-to-well contamination, cross-contamination, or sample ‘‘bleeding” as well as index switching in multiplexed sequencing libraries [37].

Contamination may affect most sequencing projects to some degree, especially low-biomass samples [38]. The composition of an environmental sample is mostly unknown before sequencing, increasing the complexity of detecting contamination when compared to a defined isolate genome and targeted sequencing project. Low-biomass samples (e.g. meconium, blood, human tissues) yield little to no DNA to be amplified and sequenced, an ideal scenario for exogenous contaminants to out-compete and dominate the biological signal.

It is important that contamination is acknowledged, accounted for and discovered at the earliest stage of a study prior to statistical analysis, to not bias measurements and to ensure that bias is not propagated into databases [39, 40]. Inclusion of negative and positive control samples is the recommended way to measure, detect, and mitigate contamination [2, 38, 41]. Negative controls should be included in the study design for every sample, extraction or amplification batch. Once provided, controls should be carefully analyzed *in-silico* and results obtained should be applied to biological samples in terms of prevalence (e.g. observations in negative controls) but also base on the frequency in relation to DNA concentration [42, 43].

However, due to the complexity and diverse possible sources of contamination, detection and mitigation are not a trivial tasks. Several approaches to identify and exclude background contamination in microbial studies have been proposed. These are based on exclusion of organisms detected in negative controls, use of replicates to find possible contaminants, removal of low abundant signals, negative correlation between organism abundance and bacterial load, clustering analysis, and others [44, 45]. Each approach has strengths and weaknesses based on the study design, data type, and control availability. Further, many studies do not include or have limited number of control samples due to the required increase in costs. [41] reported that based on publications from the 2018 issues of Microbiome and The ISME Journal, only 30% cited the use of negative controls and only 10% positive controls. Moreover, Harrison et al. [46] reported that out of 50 selected publications from 2019 and 2020, only 15 used some type of negative control and 10 of positive control to account for reagent contamination. There was also no observed increase in positive or negative controls usage in the literature from 2015 to 2020, based on selected publications. Additionally, the detection of re-occurring contaminants in extraction kits and reagents (also called “kitome”) is known to be an issue [47] but remains under-explored, mainly for not being properly cataloged, centralized or automated.

To overcome some of those challenges we propose GRIMER, a tool to analyze, visualize and explore microbiome studies with a focus on contamination detection. Based on a table of observations per sample, GRIMER generates an offline and interactive dashboard to automate data analysis, transformations and plots and generates a set of charts integrating evidence for better decision making and contamination detection. Additionally, we compiled an extensive list of common contaminants containing 210 genera and 627 species reported in 22 published articles. This data is integrated into the report. GRIMER is an effortless step once quantification is done, turning measurement tables into a interactive and dynamic report in seconds. GRIMER is open-source and available at: https://github.com/pirovc/grimer. Installation and usage instructions as well as an user manual are available in the repository. The tool is independent of analysis methods, does not rely on web or local servers and generates standalone and shareable interactive dashboards.

## Methods

GRIMER analyzes and annotates multi-sample studies based on count tables and generates a report with several interactive plots to better explore the data and to facilitate contamination detection. GRIMER integrates several sources, references, analyses as well as external tools and brings them together in one concise dashboard.

The output of GRIMER is a self-contained HTML file that can be visualized in any modern web-browser. It works independently from any actively running server or web-service. Once generated, it can be used and shared as an offline document. It has the advantages of a static report and a complex dashboard being portable and interactive. This feature makes it very convenient to distribute (e.g. as an e-mail attachment), keep track of changes in analytical pipelines and reproduce analyses in different environments.

GRIMER is independent of any quantification method and only requires a contingency table with raw counts of observations/components for each sample/composition in the study. Observations are usually, but not limited to, taxonomic entries (e.g. genus, species, strains), operational taxonomic units (OTUs), amplicon sequence variants (ASVs), or sequence features. A count of unclassified or unassigned observations is also supported to generate normalized values. Additional files and data can be provided to expand GRIMER reports: study metadata, a taxonomy database, multiple control samples, the DNA concentration, custom contaminants, and reference groups of interest. The more information provided, the more complete and interactive the final report will be.

### Annotation

GRIMER annotates observations and samples linking data with external data sources.

Sample annotations are based on a user-provided study metadata, where each sample is described in one ore more fields and variables. Those fields can contain either numeric or categorical values and are useful for grouping and clustering analyses as well as detection of batches and control/treatment effects.

Observation annotations are based on external lists of taxonomic entries, which can be used, for example, to link findings to common contaminants or connect analyses outcomes with known environments or biomes. Those entries can be easily provided by the user in a simple list of names or taxonomic identifiers in a formatted and annotated file (more information can be found in the GRIMER repository).

#### Contamination references

We compiled an extensive list of possible contaminant taxa reported in several studies (Table 3). The studies selected were obtained from cross-references in review articles [38] and individual selected findings in the literature, usually focusing on contamination detection or mitigation. Articles were manually curated and more studies can potentially be added to the list, which is dynamically maintained. Contributions are welcome through the GRIMER repository (https://github.com/pirovc/grimer/). The studies selected are very diverse in terms of sequencing technology, methodology used and environment studied. Contamination in those studies can originate from diverse sequencing kits and reagents as well as the lab environment or other unknown sources. The idea behind compiling this list is to detect which taxa is the most recurrently identified as contaminant in diverse conditions, providing a guideline and consensus for further studies. Entries on this list are not strictly considered a contaminant and should not be used alone to define contamination in a study. However, it serves as an additional evidence supporting it, especially if entries are highly recurrent (Table 4) and corroborate with additional lines of evidence. Those contaminants were reported mainly at genus or species level in different formats, names and taxonomies. We manually curated and converted them into the NCBI taxonomy [48] nomenclature for standardized usage.

**Table 3:**
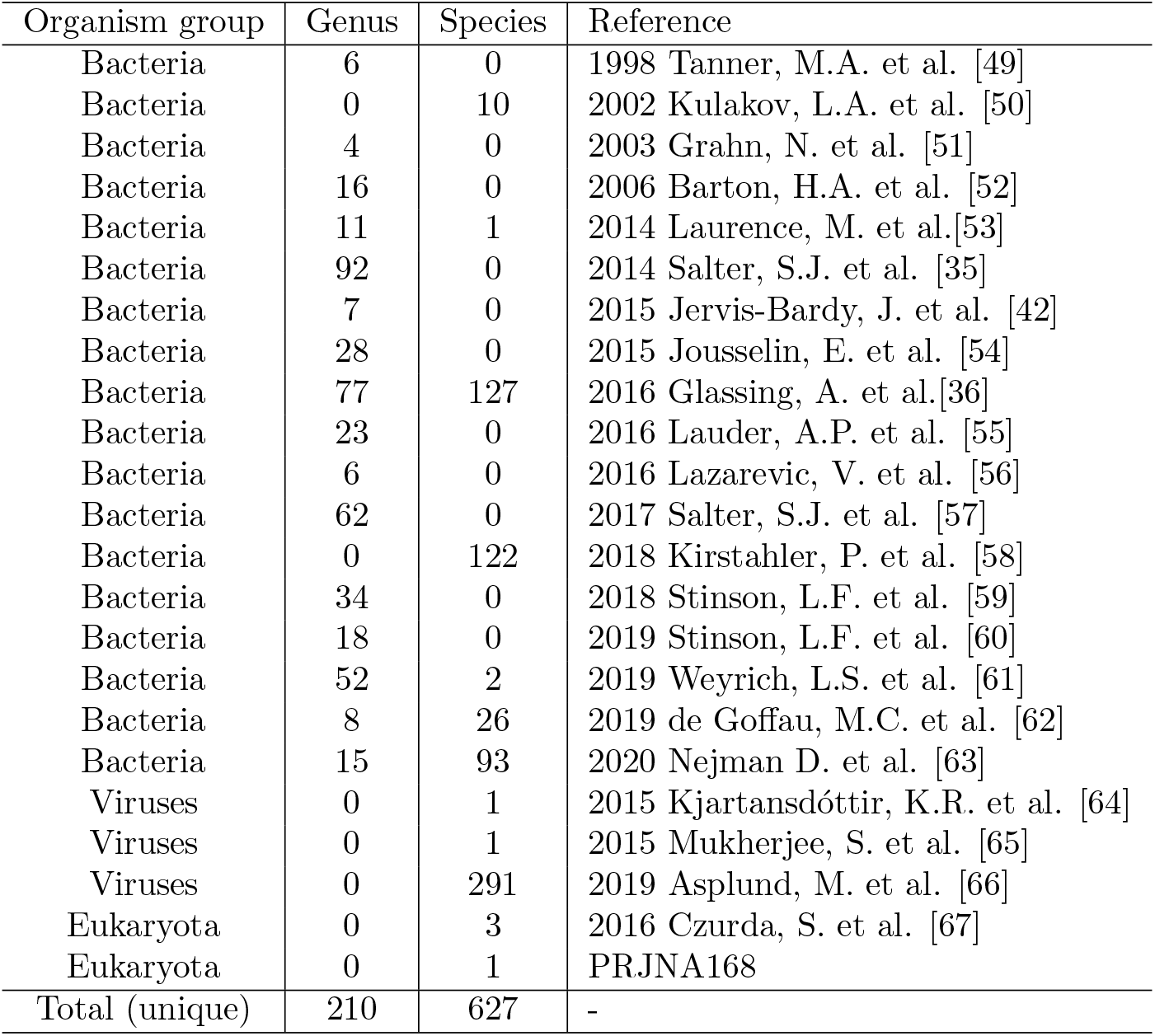
Summary of common contaminants taxa extracted from the literature. The complete list of taxa per study can be found in the GRIMER repository (https://github.com/pirovc/grimer/)

**Table 4:** Top 8 most reported taxa from Table 3 at genus and species level. If multiple child nodes of organisms are reported in the same study, they are counted here just once.

| Genus | # reported | Species | # reported |
| --- | --- | --- | --- |
| *Pseudomonas* | 13 | *Cutibacterium acnes* | 4 |
| *Stenotrophomonas* | 13 | *Pseudomonas fluorescens* | 4 |
| *Ralstonia* | 12 | *Stenotrophomonas maltophilia* | 4 |
| *Bradyrhizobium* | 11 | *Acinetobacter baumannii* | 3 |
| *Methylobacterium* | 11 | *Bradyrhizobium elkanii* | 3 |
| *Acinetobacter* | 10 | *Corynebacterium tuberculostearicum* | 3 |
| *Corynebacterium* | 10 | *Rhodococcus fascians* | 3 |
| *Sphingomonas* | 10 | *Streptococcus mitis* | 3 |

Additionally, we compiled another list of common organisms found in probable external contamination sources: taxa commonly occurring in human skin, oral and nasal cavities as well as face and other human limbs. Those were reported as possible sources of contamination [38]. Reference organisms names were obtained from BacDive [68], eHOMD [69] and further publications [70].

#### MGnify

Additionally to the contamination references, a summary generated from MGnify repository [11] is provided with counts of occurrences for each observation in thousands of microbiome studies, grouped by biome. MGnify is a resource to analyze microbiome data in an automated and standardized way. Thousands of analyzed studies are publicly available with related metadata. We mined this repository with the provided open API (https://www.ebi.ac.uk/metagenomics/api/v1/) and collected all taxonomic classifications available for every study. For each study, we collected the latest taxonomic classification based on the highest pipeline version available. If multiple classifications from different sources were present, we selected the largest one by file size. For each study output, the top 10 top most abundant organisms were linked to the study respective biome(s) definition and a final count of top organisms by biome is generated. GRIMER uses this resource to annotate observations and links how many times each identified taxa were present in other biomes. This gives another level of evidence for the possible origin of certain taxa in a study, compared to thousands of other microbiome studies. For example, in the current version, the genus *Ralstonia*, a commonly reported contaminant, appeared in 30 Environmental Aquatic biome studies and 14 Engineered Bioreactor studies (out of a total of 79 studies) while the human-related bacterial genus *Prevotella* appears mostly in host-associated biomes (89% of occurrences). All five levels of biome classification are available for each taxonomic entry.

### Input data

GRIMER requires only a contingency table to generate the full report, either in a text/tabular format (observations and samples either in rows or columns with a header) or a BIOM file [71]. Further data can be provided to extend the report:

- Metadata: annotate samples and give further technical information. The metadata should be tabular and categorical and numerical fields are supported.
- Taxonomy: GRIMER will automatically parse a given taxonomic annotation or generate one based on the provided observations. Data will be summarized in many taxonomic levels and plots will be created accordingly. Taxonomy is fully automated for several commonly used taxonomies (NCBI, GTDB, SILVA, GreenGenes, OTT).
- Controls: one or more groups of control samples can be provided in a simple text file. Those samples will be further used to summarize data and annotate plots.
- References: custom sources of contamination or any references can be provided in addition to the precompiled ones described above.

GRIMER will parse and process the data provided and run a set of analyses:

- General data summary by observation and samples, linking references, taxonomy and metadata
- Filtering and transformation: observations and samples can be filtered to reduce noise or small counts. Transformations are applied (log, centered log-ratio, normalization) to account for the composionality of the data and improve some visualizations.
- Hierarchical clustering: one or more metrics and methods can be used to perform the clustering. The combination of all of them are executed and available in the report. For this analysis, zeros are replaced by small counts defined by the user.
- Correlation: Symmetric proportionality coefficient (rho correlation) [72, 73] is calculated for top abundant observations in the study
- DECONTAM [43]: R package with a simple method to detect contaminating taxa/observations based on two main assumptions: frequency of contaminant taxa inversely correlate with DNA concentrations and contaminant taxa are more prevalent in control samples than in biological samples. DECONTAM uses linear models based on the assumptions and frequencies of the data and outputs a score for each observation to define contamination. If DNA concentration is not provided, total counts are used instead as an indirect concentration value replacement.
- MGnify: Each taxa reported will be linked to the respective MGnify entry, reporting most common biome occurrences.

### GRIMER Report

GRIMER will generate a report/dashboard with visualizations to better understand the distribution of observation counts among samples and the connection with external annotations, metadata and taxonomy. Currently GRIMER reports contain 4 main panels: Overview (Figure 6), Samples (Figure 1), Heatmap (Figure 4), and Correlation (Figure 5). Some of them were previously suggested to be adequate for contamination detection [45] and are commonly used in standard microbiome analysis. Every panel has one or more visualization and widgets to select, filter, group, and modify its contents. Panels can be reported independently.

**Figure 1:**
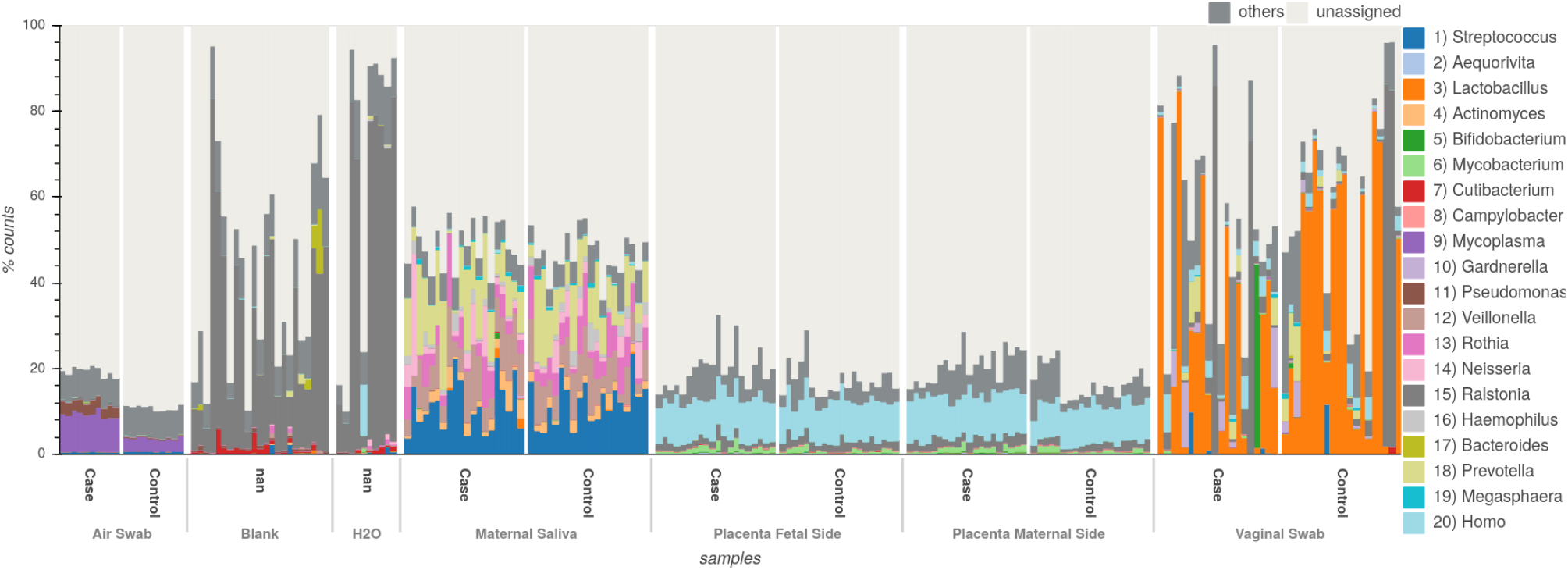
Bar plot with relative abundance of top 20 genera for the placenta study. Bars are grouped by sample type and case/control. There is a stark difference in composition between air, vaginal and saliva samples to placental samples and controls (Blank, H_2_0).

**Figure 2:**
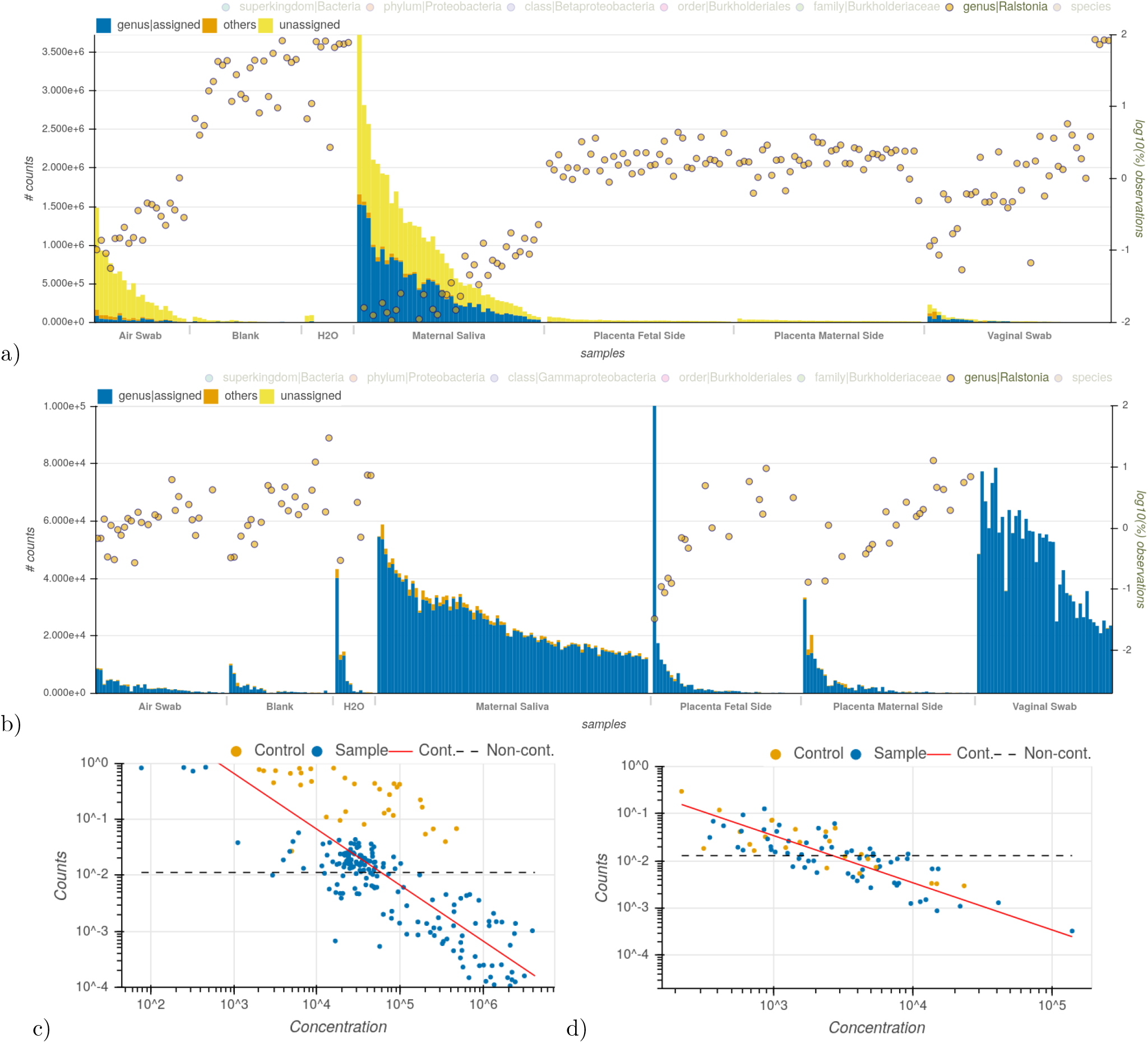
Evidence supporting *Ralstonia* as a contaminant taxa in the placenta study. a-b) Right y-axis shows normalized abundance of genus *Ralstonia* in log scale for each sample in the MGS (a) and amplicon (b) data. Bars (left y-axis) summarized counts at genus level for each sample. Samples are grouped by sample type and sorted by number of reads (x-axis). The yellow circles show abundance of *Ralstonia*, which is higher in the control samples (Blank and H_2_0) as well as increased in real samples with low read count. c-d) DECONTAM plots for *Ralstonia* genus for the MGS (c) and amplicon (d) data. DECONTAM plots show that taxa counts follow the expected distribution for contamination based on the number of reads per sample (red line).

**Figure 3:**
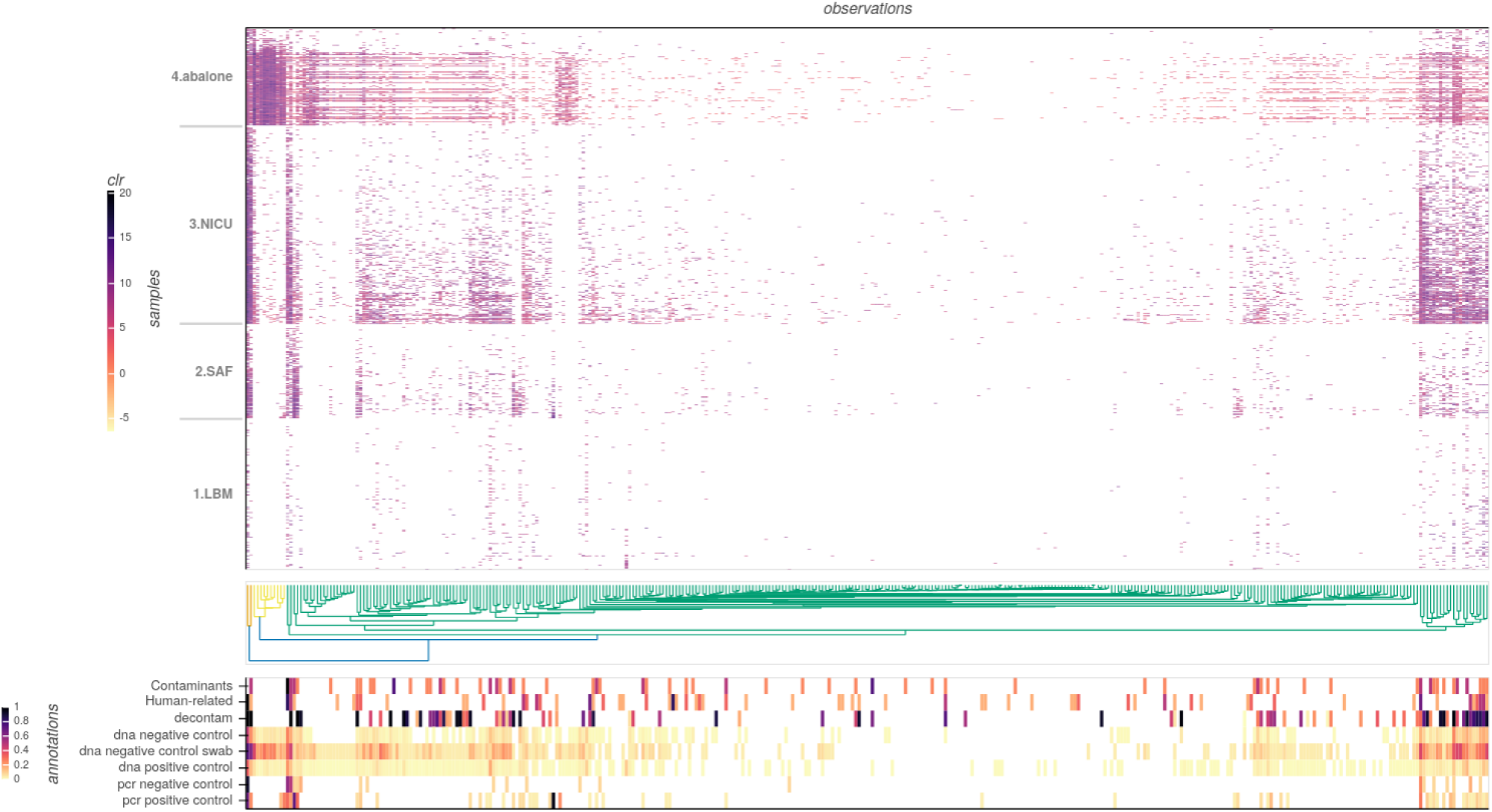
Heatmap visualization at species level for the KatharoSeq data. Samples are grouped by study type (y-axis) and clustered by observations (x-axis, euclidean distance metric, complete method). Data in the heatmap is center-log ratio transformed. Bottom panel show annotation related to the observations. ‘‘Contaminants” and “Human-related” annotations are normalized counts against pre-compiled list of references described in this paper. “decontam” is the normalized DECONTAM p-score. All “control” annotations show the proportion of the observation in the indicated group of control samples.

**Figure 4:**
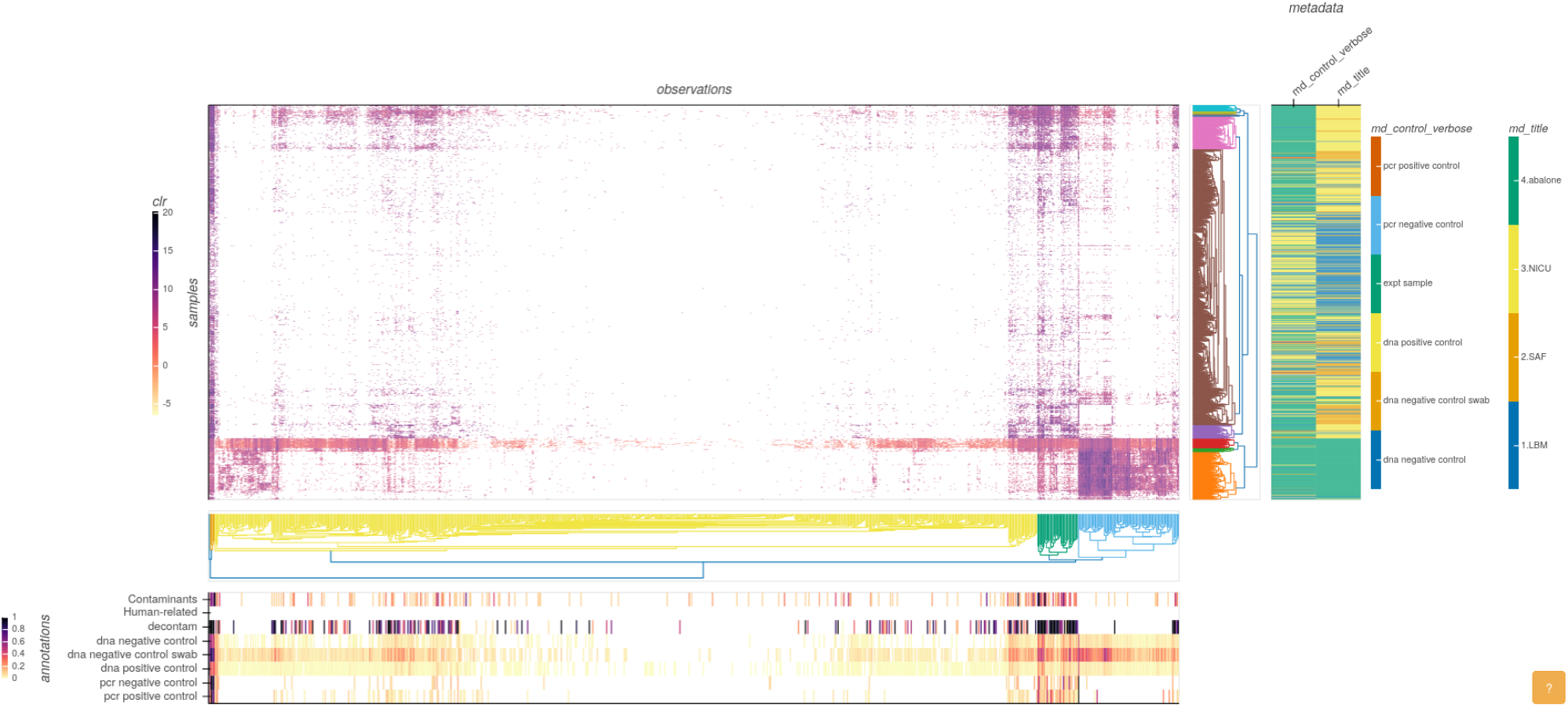
Heatmap visualization at genus level for the KatharoSeq data. Samples and observations axis are clustered and sorted based on the euclidean distance metric, complete method. Data in the heatmap is center-log ratio transformed. Bottom panel show annotation related to the observations. “Contaminants” and “Human-related” annotations are normalized counts against pre-compiled list of references described in this paper. “decontam” is the normalized DECONTAM p-score. All “control” annotations show the proportion of the observation in the indicated group of control samples. Metadata panel show color-coded sample information on study (md_title) and type of sample (md_control_verbose). The annotation panel shows higher values on multiple sources of evidence for contamination relative to data clusters of the heatmap. Metadata panel shows how samples show independent patterns based on the environment (md_title) and difference from controls (md_control_verbose).

**Figure 5:**
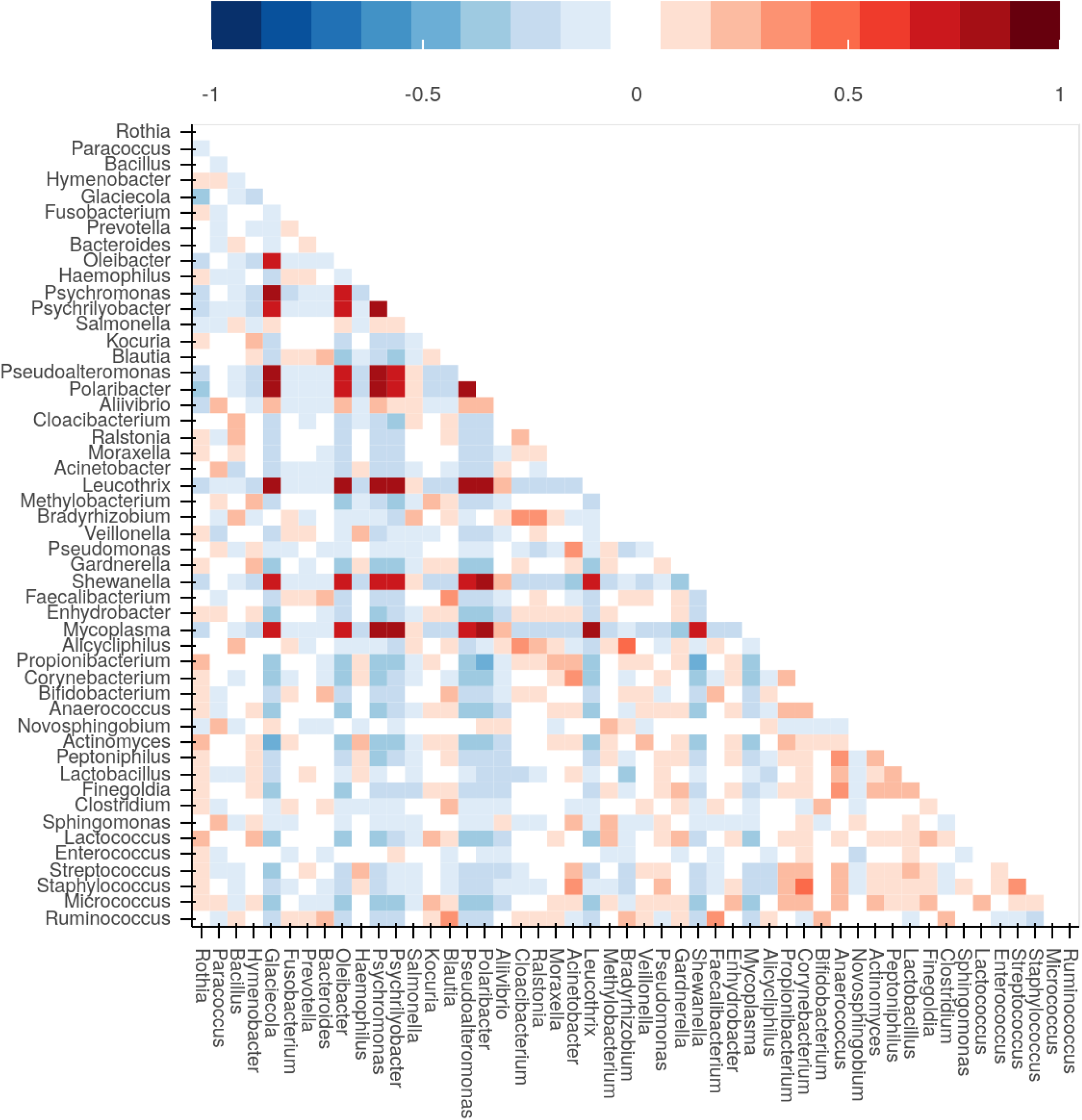
Symmetric proportionality coefficient (rho correlation) between top 50 most abundant genera in the KatharoSeq data. Positive correlation values (between 0 and 1) are displayed in red. Negative correlation values (between −1 and 0) are displayed in blue. Highly correlated matrix among 9 genera (dark red) points to reagent-derived contamination, when considered with other lines of evidence (Figure 6)

**Figure 6:**
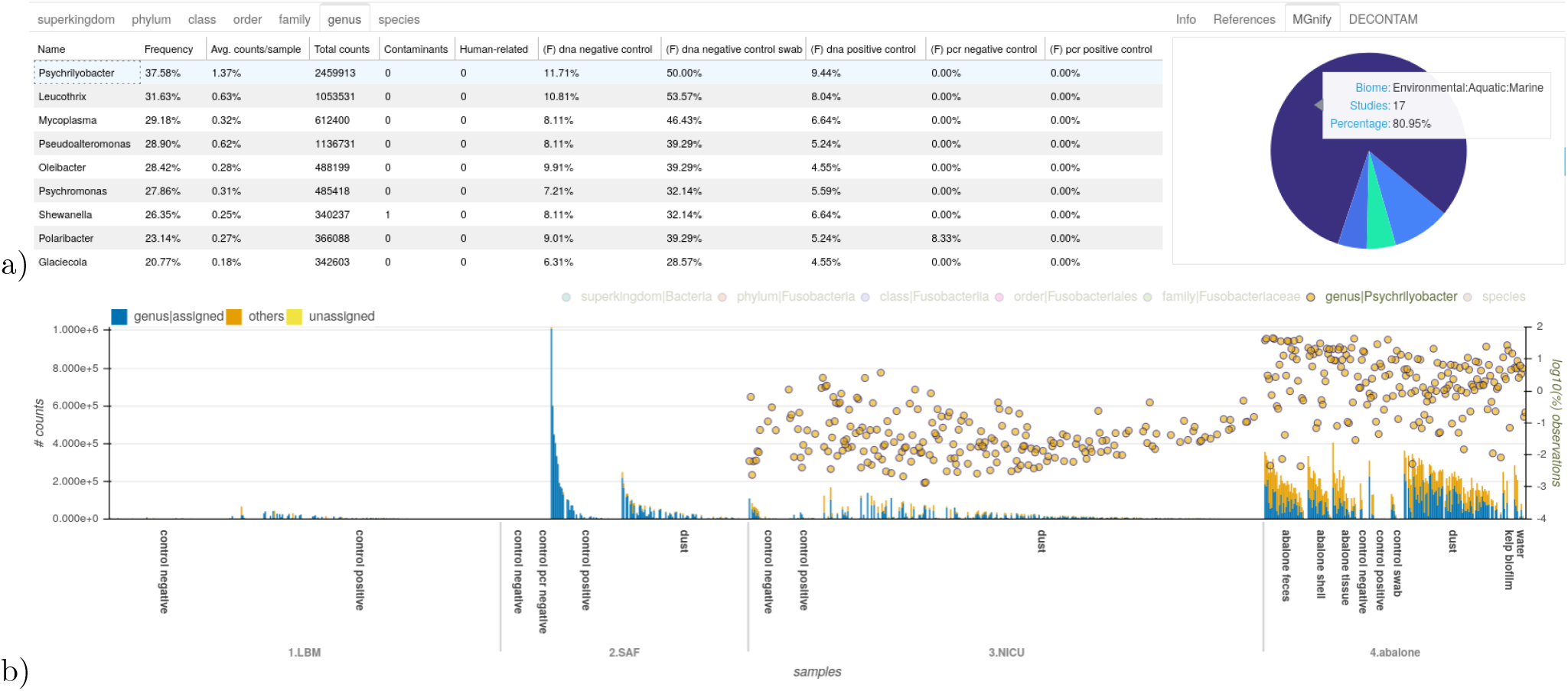
GRIMER Overview panel plots a) listing of 9 highly correlated genera detected in Figure 5. Samples have high incidence in DNA negative controls. MGnify plot showing proportion of biomes related to *Psychrilyobacter* in the whole MGnify database b) bar plot listing samples (x-axis), grouped by study and sample type and sorted by total number of reads. Bars represent the total number of counts for each sample and are annotated with the proportion assigned to genus level (left y-axis). Log-transformed abundance of *Psychrilyobacter* is displayed in yellow circles (right y-axis). This taxon is abundant in the abalone samples but has some signal in the NICU samples that are inversely correlated to the total amount of reads, pointing to potential contamination. The other 8 taxa from show similar patterns in the report.

An individual summary for each observation and its relation to annotations and distribution among samples can be found in the Overview panel (Figure 6). Here, all evidence related to a specific observation are integrated for further examination. Each observation provided in the study is listed and summarized in a tabular format. Once selected, the distribution of counts for the specific observation for each sample can be observed in a bar plot. Information of annotations, MGnify biomes, and DECONTAM output are also available in the same interface. The DECONTAM output indicates if the observation is classified as a contaminant with a score and a plot showing the frequency of the selected observation against the DNA concentration for all samples containing that observation. Linear models showing the expected values for contamination and non-contamination values are also plotted. If provided, taxonomic lineages are integrated in the table and plots and observation are decomposed and summarized into taxonomic levels. The Overview panel also roughly summarizes samples contents in the bar plot, with general classification metrics. Those bars can be transformed, annotated, grouped and sorted to connect specific observation values to overall sample distribution.

In depth evaluation of individual samples can be performed in the Samples panel (Figure 1). Normalized distribution of top observations for each sample can be visualized in the bar plot to easily compare the overall distribution of observations among samples, with options for grouping and sorting by metadata. Automated selection of groups of samples is also possible by counts and metadata.

Several transformations can be applied to the data (normalization, log, center log ratio) to be further visualized in the Heatmap panel (Figure 4). Hierarchical clustering, grouping and sorting options can be independently selected for samples and observations to enable pattern detection (e.g. batch effects, treatment effects etc). Dendrograms are plotted when clustering options are selected. Annotation bars are plotted around the heatmap showing dynamically selection of sample annotations (metadata) and observation annotations (references, controls and DECONTAM output). Metadata is automatically colored to reflect categories (distinct colors) and numeric (sequential colors) fields. Multiple metadata fields can be select interactively. Observation annotation values are normalized and plotted in the same color scale for easier interpretation. One heatmap is generated for each taxonomic level.

Correlation between observations are plotted as a matrix (Figure 5). Positive or negative correlations among observations can point towards concurrent signals in the microbiome analysis. Observations present in multiple samples in similar ratios are positively correlated and the opposite configures negative correlation. Once a signal is observed, the correlation matrix can indicate co-occurrence of observations and help to identify further candidates (e.g. cluster of co-occurring contaminants at similar ratios).

## Implementation

GRIMER is written in Python and Javascript and outputs a report file in HTML format. All visualizations and layouts are created with the the Bokeh library (https://bokeh.org). Bokeh plots, tables and charts automatically provide a set of tools for interaction (e.g. zoom, selection) with an option to export the current selection to an image file. Many plots have interactive tool-tips, showing more information about the data under the mouse cursor. Help buttons are also included, explaining the plots and options.

Further libraries were used to analyze samples and generate the report: pandas [74] for general parsing and data structures, scipy [75] for hierarchical clustering, scikit-bio (http://scikit-bio.org/) for transformations. Scripts to download and generate MGnify annotations and update reference sources are provided in the GRIMER repository (https://github.com/pirovc/grimer).

GRIMER automatically handles taxonomic entries using MultiTax (https://github.com/pirovc/multitax). GRIMER will automatically parse given taxonomies or download and convert any taxonomic id or name internally and decompose results in taxonomic ranks. Currently supported taxonomies are: NCBI, GTDB, Silva, GreenGenes, OpenTree Taxonomy. Reference lists are currently only available based on the NCBI Taxonomy.

## Results

We re-analyzed publicly available studies to demonstrate the use of GRIMER reports in real case scenarios and what types of analyses are possible. In some examples, we try to reproduce analyses and in other cases point to new evidence that may have been overlooked. We encourage the readers to open the GRIMER reports (https://doi.org/10.5281/zenodo.7525798 or https://pirovc.github.io/grimer-reports/) and interactively visualize the results being described to fully understand the capabilities of the report. All reports presented below were generated using GRIMER version 1.1.0.

### Detecting contamination

The attempt to detect and describe a possible human placental microbiome has motivated several studies and investigations [76, 47, 62, 77]. Leiby et al. [78] published a detailed and well designed study contributing to the subject. Placental samples for term (control) and preterm (case) newborns were collected for the maternal and fetal sides. Additionally, positive control samples were obtained from the mothers (saliva and cervicovaginal fluid) as well as negative control samples (air from the sample processing room, empty tubes and PCR grade water). The study was performed in both marker gene sequencing (amplicon) and metagenomics (MGS). The authors could not distinguished a unique placental microbiome which differs from the contamination background. We re-analyzed the samples in a standard pipeline with QIIME2 [6] for amplicon data and ganon for MGS data [79], generated a GRIMER report for both and searched for the previously detected contamination.

In the MGS report, the bar plot (Figure 1) shows a stark difference in signal between sample types but a smaller difference in case and control groups. The *Ralstonia* genus is present in 96% of the all samples with an average abundance of 8.24%. Reads assigned for this genus were found in all negative control samples and H_2_0 samples. *Ralstonia* was also reported in 12 studies as a common contaminant, based on our compiled contaminant list (Table 4) and it was classified as a contaminant by the DECONTAM method, based on the correlation of frequencies and the total number of reads per sample. Further, the abundance of this genus is higher in negative controls and placental samples as well as in samples with low number of reads, probably related to their low biomass as depicted in the Figure 2. Those results are inline with the ones reported in the original publication [78], even though the data was re-analyzed with a different set of tools, parameters and reference databases. All evidence describe pointing to *Ralstonia* as a contaminant was automatically generated by GRIMER and can be directly extracted from the Overview panel from the report. Besides human reads, *Ralstonia insidiosa* is the most prevalent species in this study. For the amplicon data, a similar pattern can be detected for the *Ralstonia* genus based on amplicon sequence variants (Figure 2).

Further, all other taxa present can easily be verified for the same patterns. *Pseudonomas* show similar distribution and was also reported originally as probable contaminants in the placental samples. *Corynebacterium, Cutibacterium* and *Mycobacterium*, although less prevalent, are further taxa with very similar patterns that could be potential contaminants and were not reported in the original publication.

### Multiple microbiome studies exploration

Definitive and robust conclusions from low-biomass studied environment are only possible with a set of controls and protocols to deal with contamination. KatharoSeq [80] is a well designed protocol to better handle contamination in high-throughput low-biomass DNA microbial studies for amplicon sequencing or shotgun metagenomics. The protocol has guidelines for positive and negative controls implementation at the DNA extraction and library construction steps as well as computational approaches to define and exclude samples that did not achieve minimal amount of signal to be used. In their publication [80], the authors validate the protocol sequencing and analyzing with 3 low-biomass environments: the Jet Propulsion Laboratory spacecraft assembly facility (SAF), rooms of a neonatal intensive care unit (NICU) and an endangered-abalone-rearing facility (abalone). A set of negative controls to compare extraction kits is also included in the study (LBM).

We downloaded the OTU table and metadata from KatharoSeq evaluations for the 16S rRNA analyses available in Qiita [14] in the following configuration: reads trimmed at 150bp and classified using closed-reference OTUs clustered at 97% similarity annotated with the greengenes taxonomy. A GRIMER report was generated for the raw table with all samples without any filtration. The heatmap generated for the annotated species level (Figure 3) shows a distinct and clear pattern between environments and the LBM. As reported in the publication, abalone samples have a higher richness (here as species annotated OTUs) as well as the highest average number of reads per sample. It is possible to identify potential contaminants in the study by looking for observations prevalent across environments and the relation to its annotations. Using this analysis we detected *Cutibacterium acnes* which is reported as a common contaminant and human-related species, present among all 4 environments studied as well as highly frequent in negative and positive controls. Even though DECONTAM did not identify this taxa as a contaminant, related data still hold strong evidence for contaminant of *C. acnes* in this study. Furthermore, *Staphylococcus aureus* and *Staphylococcus epidermidis*, known as human-related bacteria, were detected in high abundances in both NICU and SAF environments - areas with low and high human exposure, respectively. However, both species were also relatively highly present in negative controls, the abalone environment and LBM samples. Additionally, both were positively classified as contamination by DECONTAM, indicating that besides human exposure, those organisms could be driven by an external source of contamination.

Species identification based on 16S rRNA is limited due to its low resolution: approximately 15% of the OTUs are annotated at species level and 69% at genus level in this study. The same analysis visualized at genus level gives an increased perception of the distribution of the data in this study. With a higher signal it is possible to visualize how several clusters are formed and in many cases agree in multiple levels of evidence supporting the possibility of contamination (Figure 4).

Looking at the correlation between top observations reported (Figure 5), a matrix of highly correlated genera can be detected. Such a pattern was previously reported to be an indication of contamination from reagent-derived sources since they are invariably present within samples in similar ratios [45]. Further inspection of those genera (*Glaciecola*, *Leucothrix*, *Mycoplasma*, *Oleibacter*, *Polaribacter*, *Pseudoalteromonas*, *Psychrilyobacter*, *Psychromonas*, *Shewanella*) shows that they are mainly from Aquatic/Marine biomes with help of the matching results with the MGnify database (Figure 6 B). Further, they are more prevalent in negative controls (Figure 6 A), an evidence of DNA extraction kit or sample processing contaminants. Those organisms are highly frequent in the abalone study, which is a Marine environment and some of them were also described in the original publication. Although in very low amounts, those groups were also reported present in NICU samples (Figure 6 C), pointing to possible well-to-well contamination.

## Discussion

GRIMER is an easy-to-use and accessible tool for specialists and non-specialists that generates a concise interactive offline dashboard with a set of analyses, visualizations, and data connections from a simple table of counts. It automatically summarizes several levels of evidence to better understand the relation between observations, samples, metadata, and taxonomy. GRIMER reports are a valuable resource for investigating contamination, a problem that affects every microbiome study to some degree.

All the conclusion and visualizations presented in this work in the results section were solely based on GRIMER reports, showing that microbiome analysis, contamination investigation and detection are possible with the methodology proposed. The use of multiple sources of evidence to annotate observations improves the ability to better detect clear contaminants in microbiome studies as well as to point to probable groups of candidate contaminants.

In addition to the GRIMER software, we compiled and provided in this work a list of common taxa contaminants based on 22 publications (Table 3). Many of the reported contaminants are recurrent in diverse studies, pointing to a consensus for some taxa (Table 4) as a probable contaminant. Taxa in this list cannot be strictly considered a contaminant by itself. However, they can corroborate suspicious contamination discovered via several other lines of evidence without the extra effort of researching the literature. The presented list is not comprehensive but a first step to centralize and standardize re-occurring contaminants described in the literature. We expect this list to incrementally grow overtime as more evidence of kit and laboratory contamination becomes available. The information of common contaminants is a valuable resource to aid contamination detecting and we are willing to keep and extend it. Improvements to the list and suggestions of further candidate taxa can be provided via the GRIMER repository at https://github.com/pirovc/grimer/. As a future work, the list can be associated with study details as biome, extraction kit and methodology to be further queried and integrated in more details.

Additionally to the aforementioned common contaminants, GRIMER can also use general lists of custom organisms to annotate samples. In this work and by default, human-related organisms commonly occurring in human skin, oral and nasal cavities as well as face and other human limbs are used since they can be external sources of contamination. Those lists can be easily provided as taxonomic identifiers or names to GRIMER. If the target study conflicts with any of those environments (e.g. study of human skin), one could simply remove the related entries from the configuration files. More details and examples on how to perform this can be found in the online documentation.

GRIMER works out-of-the-box with as little data as possible but can incrementally expand the reports when more data is provided and can be adapted for user necessities. GRIMER is fast and generate reports in a matter of seconds on a standard notebook. The outcome dashboard is lightweight and can handle hundreds to thousands of samples and observations. Report sizes usually vary from 1-10MB and are highly compressible, since they are text-based HTML files. GRIMER reports with higher number of samples (thousands) can grow significantly in size (10-100MB) but still run normally. If report size is a limitation, many options can be adjusted to generated more compact files: reducing number of taxonomy ranks displayed, less combinations of analyses, filtering very low abundant observations, among others.

One of GRIMER core strengths is the taxonomy automation. It accepts taxonomic identifiers from several different taxonomies, but also parses names and converts them to their respective identifiers. If only one taxonomic level is provided (e.g. species level), GRIMER can decompose and summarize the data in higher ranks. That means that users do not have to handle taxonomy and everything will work automatically. GRIMER was developed in a way that new visualizations can be included with little effort.

We listed and summarized a list of similar currently available methods published in the last 10 years (Table 2) as well as web-plataforms for complete analyses of microbiome data (Table 1). A list of functionalities between similar available tools is provided in [9] but a detailed comparison with GRIMER is out of the scope of this work. Most methods share some basic functions (e.g. taxonomic abundance analysis) but are diverse in many other aspects and were sometimes developed with specific goals (e.g. function analysis, biomarker identification). However, there is no comprehensive method that can provide a complete solution for the many possible analyses in a microbiome study. We believe that many of those tools, besides their overlapping functions, are complementary and can be used concurrently. GRIMER mainly shares features with pavian [22] in terms of general microbiome exploration and support to metagenomics data and with OpenContami [28] regarding contaminant detection. GRIMER, however, is unique in its output format. The vast majority of the currently available tools are web-based, hosted in a remote server or rely on a local hosted web-server to properly work (Table 2). This may be impractical for many non-specialists and for long term storage and reproducibility. GRIMER reports are portable and fully functional offline. This allows analysis to be accessible by many researchers with different backgrounds working together in the same study, increasing direct interaction with data. The portability also enables better documentation of results, reproducibility and shareability. Further, web-based tools may disappear after some years of inactivity or lack of funding and analysis may be lost, as it is the case for for some methods (Table 5). GRIMER reports are completely offline and will work as long as the report file is safely stored.

**Table 5:** Tools and web resources no longer available, supported or inaccessible (as of 2022-02-28)

| Name | Reason | Year | Reference |
| --- | --- | --- | --- |
| Community-analyzer | website offline | 2013 | [81] |
| calypso | website offline | 2017 | [82] |
| Metaviz | web tool not responsive | 2018 | [83] |
| iMAP | no longer supported due to funding | 2019 | [84] |
| biomminer | page not found | 2020 | [85] |

Overall we believe that GRIMER is a valuable contribution to the microbiome field and can facilitate data exploration, analysis and contamination detection.

## Declarations

### Availability of data and materials

The datasets and metadata for the placenta study were obtained from: https://www.ebi.ac.uk/ena/browser/view/PRJNA451186

The datasets and metadata for the KatharoSeq study were obtained from (log-in required): https://qiita.ucsd.edu/study/description/10934

### Authors’ contributions

V.C.P and B.Y.R. conceptualized and developed the idea. V.C.P. wrote the software and main manuscript text, including all analyses. B.Y.R. reviewed and contributed to the manuscript text. All authors reviewed the manuscript.

## Acknowledgements

We thank all partners in the ZooSeq project for helpful discussions and support in this project.

## Availability and requirements

Project name: GRIMER

Project home page: https://github.com/pirovc/grimer

Operating system(s): Platform independent

Programming language: Python 3.5 or higher

Other requirements: bokeh 2.2.3 or higher

License: MIT License

Any restrictions to use by non-academics: use based on MIT licence

